# Genetic behavior analysis for phytochemical traits in coriander: Heterosis, inbreeding depression and genetic effects

**DOI:** 10.1101/2021.09.23.461492

**Authors:** Amir Gholizadeh, Mostafa Khodadadi

## Abstract

Increasing fruit yield, fatty acids and essential oils content in coriander are the main objectives. Reaching them need to understand the nature of gene action and quantifying the heterosis and inbreeding depression. Six genetically diverse parents, their 15 F_1_ one-way hybrids and 15 F_2_ populations were evaluated under different levels of water treatments. Beside the water treatment and genotype effects, the genetic effects of general (GCA) and specific (SCA) combining ability and their interactions with water treatment were significant for all traits. Water deficit stress decreased all traits in both F_1_ and F_2_ generations except for essential oil content which were significantly increased due to water deficit stress. Under water deficit stress, a non-additive gene action nature was predominant in F_1_ generation while an additive gene action nature was more important in F_2_ generation for all the traits except fruit yield under severe water deficit stress. There was a positive high heterosis for the traits examined in some hybrids. Also, in F_2_ generation even after inbreeding depression, some promising populations displayed appropriate mean performance. These show that the parents used for crossing had rich gene pool for studied traits. Therefore, selection between the individuals of relevant F_2_ populations could be led to develop high yielding hybrids or transgressed lines.

## Introduction

Coriander (*Coriandrum sativum* L.) is a member of Apiaceae family which has known as medicinal and industrial plant. Food characteristics caused to cultivate and wide spread of coriander. It is used for different applications such as food, drugs, cosmetics and perfumery industry (Neffati and Marzouk, 2008). Coriander fruit contains both fatty acids and essential oils. A petroselinic acid is a main component of the fatty acid consisting 85% of the total fatty acids. In industry, petroselinic acid is broken-down into lauric, adipic and C_6_ dicarboxylic acids which are used for synthesizing detergents and nylon polimer (Murphy et al., 1994; Murphy, 1996). The fatty oil composition of coriander fruit has previously been characterized (Ramadan and Morsel, 2002; Ramadan and Morsel, 2006; Msaada et al., 2009a; Sriti et al., 2009).

The essential oils in coriander have become interesting alternative for other natural components in food (Wong and Kitts, 2006; Donega et al., 2013). Also, the essential oils are used to flavor or remove unpleasant odors of some products in food industry (Matasyoh et al., 2009; Neffati and Marzouk, 2010). Essential oil composition of coriander fruit has previously been quantified (Msaada et al., 2007; Msaada et al., 2009b; Sriti et al., 2009, Neffati et al., 2011). Coriander essential oil includes 60-70% linalool has the pleasant characteristics odor (Lubbe and Verpoorte, 2011). Also, many medicinal properties have been attributed to coriander essential oil, including antibacterial (Burt, 2004; Lo Cantore et al., 2004), antioxidant (Wangensteen et al., 2004), antidiabetic (Gallagher et al., 2003) and anticancer (Chithra and Leelamma, 2000) and anti-antimicrobial activities (Matasyoh et al., 2009; Begnami et al., 2010; Neffati et al., 2011).

It was revealed that the amount and composition of substances and secondary metabolites affected by water deficit stress in some medicinal plants (Charles et al., 1990; Petropoulos et al., 2008). In some studies, an enhancing effect of water deficit stress on the biosynthesis of essential oils observed (Jaafar et al., 2012; Alinian et al., 2016). Under the stressful growth condition, secondary metabolites and/or substances production in plants enhanced for preventing an oxidization in the plant cells. Similarly, under water deficit stress an essential oil content may be increased. In case of fatty acids, there are evidences about the decreasing effect of water deficit stress on fatty acids content and yield (Hamrouni et al., 2001; Bettaieb et al., 2009; Bettaieb et al., 2011). To decrease adverse effects of drought stress on farmers’ economy through lowering the yield of common crops, cultivation of medicinal plants with improved potential of secondary metabolites production under drought-affected areas could be suggested as an alternative approach (Alinian et al., 2016).

Consideration the statements, for increasing essential oil and fatty acids yield in coriander, reaching drought-tolerant cultivars with high fruit yield and fatty acids and essential oil content through plant breeding could be possible. Generally, plant breeding is known as a more stable approach and a complementary for decreasing the deleterious effects of water deficit stress through the development of genotypes which can grow and produce suitable essential oil yield under water deficit stressed environments. Any successful plant improving program depends on an understanding the nature of gene action involved in the inheritance of that traits under target growth condition. Griffing’s (1956) diallel analysis has used to uncover the behavior of genes involved in controlling of the traits. This method has also used to estimate variance of GCA and SCA in different self-pollinated and open-pollinated crops (Khan et al., 2009; Blank et al., 2012; Townsend et al., 2013; El-Gabry et al., 2014; Khodadadi et al., 2016b; Khodadadi et al., 2017; Kaushik et al., 2018; Teodoro et al., 2019; Schegoscheski Gerhardt et al., 2019).

The heterosis phenomenon in F_1_ hybrids can address the SCA and GCA of relevant parents. Therefore, heterotic breeding search for valuable hybrid combinations which have the commercialization potential. On the other hand, inbreeding depression measures the amount of vigor reduction in segregating generations due to self-pollination (Joseph and Santhoshkumar, 2000).

Diallel analysis on F_1_ crosses has previously been done to estimate genetic parameters and combining ability in coriander (Khodadadi et al., 2016b). But, it is necessary to uncover the heterosis, inbreeding depression and repeatability of genetic estimates through F_2_ diallel analysis to establish a successful breeding program for improving coriander fruit quantity and quality under water limiting conditions in coriander. The objectives of this study were understanding gene action nature in controlling fruit yield and some phytochemical traits and identifying heterosis and inbreeding depression potential in coriander under different levels of water treatment.

## Materials and methods

### Plant material and growth conditions

Genotypes used for making diallel crosses had been evaluated in a preliminary experiment for drought tolerance by Khodadadi et al. (2016a). The characteristics of selected parental genotypes were summarized in Table 1. All the six parents contributed to produce 15 F_1_ hybrids (without reciprocals) through half diallel mating system in 2015. A part of these F_1_ hybrids’ seed were used to produce 15 F_2_ generations through self-pollination in the isolated condition. All of the six parents, 15 F_1_ hybrids and 15 F_2_ generations were evaluated under three levels of irrigation regimes. A field trial consisted three experiments close together 1 meter distance. These experiments were well watered (WW), moderate water deficit stress (MWDS) and severe water deficit stress (SWDS). Each of these experiments carried out through the randomized complete block design with three replications at the research field of Tarbiat Modares University (51° 09 ′E; 35° 44′ N; altitude 1265 m), Iran during the growing season of 2017. In WW experiment, a set of genotypes were well watered overall the experiment period. In MWDS experiment, a set of genotypes were well watered until an appearance of the stem when watering was withdrawn until the end of the flowering stage at which point one recovery watering applied. In SWDS experiment, watering was similar to WW experiment until an appearance of flowering stage and after which watering was cut off completely. The research field soil physical and chemical characteristic presented in Table 2.

**Table 1.**
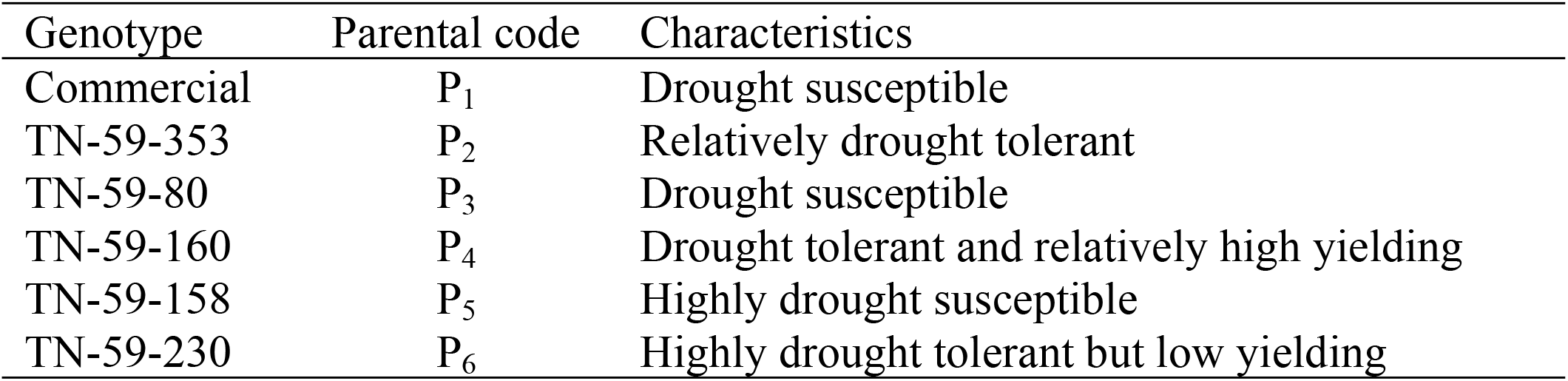
Coriander genotypes and their characteristics.

| Genotype | Parental code | Characteristics |
| --- | --- | --- |
| Commercial | P<sub>1</sub> | Drought susceptible |
| TN-59-353 | P<sub>2</sub> | Relatively drought tolerant |
| TN-59-80 | P<sub>3</sub> | Drought susceptible |
| TN-59-160 | P<sub>4</sub> | Drought tolerant and relatively high yielding |
| TN-59-158 | P<sub>5</sub> | Highly drought susceptible |
| TN-59-230 | P<sub>6</sub> | Highly drought tolerant but low yielding |

**Table 2.** Soil properties of different layers of the experimental field.

| Soil depth (cm) | Sand (%) | Silt (%) | Clay (%) | Bulk density (g cm<sup>-3</sup>) | FC (%) | Organic matter (%) | pH | EC (dS m<sup>-1</sup>) |
| --- | --- | --- | --- | --- | --- | --- | --- | --- |
| 0-20 | 70 | 15 | 15 | 1.2 | 16.5 | 1.61 | 7.75 | 1.3 |
| 20-40 | 68 | 18 | 14 | 1.4 | 19 | 1.45 | 7.75 | 1.28 |
| 40-60 | 66 | 18 | 16 | 1.48 | 15 | 1.09 | 7.74 | 1.26 |

FC, soil moisture at field capacity.

### Trait Measurements

The phytochemical traits include essential oil content (EOC), fatty acid content (FAC), essential oil yield (EOY) and fatty acid yield (FAY), fruit yield per plant (FY) were measured. For measuring fruit yield of parents and relevant F_1_ hybrids 10 plants were harvested from each of the experimental plots. In F_2_ generations 30 plants were harvested from each of the experimental plots. For extracting the essential oil, 30 g of dried coriander fruits were well powdered and subjected to hydro-distillation in Clevenger-type apparatus for 120 min. Essential oil content (%w/w) was computed through the weight (g) of essential oil per 100 g of fruit (Khodadadi et al., 2016b). Also, essential oil yield was computed through multiplying the essential oil content by fruit yield per plant (g). For measuring fatty acid content, two grams of powdered fruit sample of coriander were subjected to Soxhlet apparatus with 250 ml of petroleum ether for 6 h. Fatty acids were removed after mixture filtration and solvent evaporation under reduced temperature and pressure (Alinian and Razmjoo, 2014; Khodadadi et al., 2016b). Finally, fatty acid yield was estimated by multiplying fatty acid content with fruit yield per plant (g) for each plot.

### Statistical analysis

The datasets were firstly tested for normality using the Anderson and Darling normality test. The analysis of variance for GCA and SCA effects were done according to Griffing’s (1956) method 2, model 1 using a SAS program suggested by Zhang et al. (2005). Mean values of traits in water treatments were compared using the least significant difference (LSD) method at 5% level of probability. Estimates of 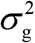 (general combining ability variance) and 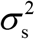 (specific combining ability variance) were computed according to the random-effects model (Zhang et al., 2005). The GCA /SCA ratio was computed according to the method proposed by Baker (1978) (Equation 1).

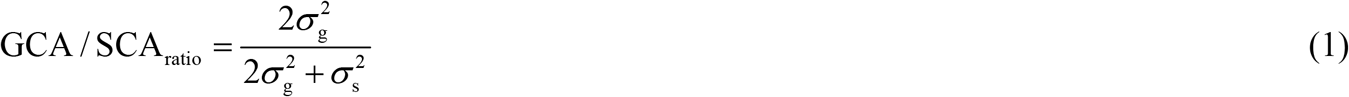

The best parent heterosis was calculated in F_1_ hybrids using the formula suggested by Fonseca and Patterson (1968) (Equation 2).

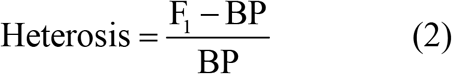

where F_1_ and BP are target hybrid and best parent values, respectively. Also, the observed inbreeding depression (ID) was estimated as a percent of the decrease in F_2_ mean when compared with F_1_ hybrid mean according to the formula suggested by Khan et al. (2009) (Equation 3). The 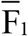 is the mean value of F_1_ hybrid and 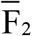 is the mean value of F_2_ generations mean of parents.

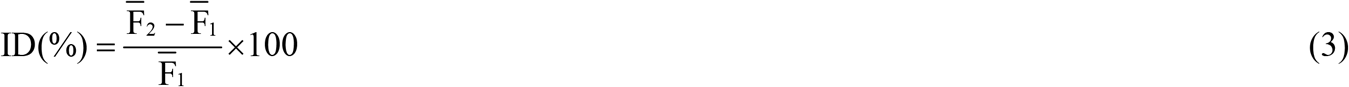

All statistical analysis were done using Statistical Analysis System (SAS) (SAS Institute, 1992) and graphs generated using Excel Microsoft Office Software.

## Results and discussion

### Combined analysis of variance for traits under water treatments

The combined analysis of variance revealed the presence of a significant difference between water treatments for all of traits in both F_1_ hybrids and F_2_ generations (Table 3). There was a high significant difference between F_1_ hybrids and also between F_2_ generations for all of studied traits. These observations indicate that parent selection for diallel crosses had been properly done. Along with the main water treatment and genotype effects, the genotype × water treatment interaction effect was significant for all traits in both F_1_ hybrids and F_2_ generations (Table 3). Being significant genotype (F_1_ hybrids + F_2_ generations) × water treatment interaction refers to different growth response of genotypes in differently watered growth conditions.

**Table 3.** Combined analysis of variance for phytochemical traits in the F_1_ and F_2_ generations under water treatments

| Source | df | Mean Squares |  |  |  |  |  |  |  |  |  |
| --- | --- | --- | --- | --- | --- | --- | --- | --- | --- | --- | --- |
|  |  | FY |  | EOC |  | FOC |  | EOY |  | FOY |  |
|  |  | F<sub>1</sub> | F<sub>2</sub> | F<sub>1</sub> | F<sub>2</sub> | F<sub>1</sub> | F<sub>2</sub> | F<sub>1</sub> | F<sub>2</sub> | F<sub>1</sub> | F<sub>2</sub> |
| Water treatment (WT) | 2 | 771.31** | 332.34** | 0.53** | 0.193** | 223.12** | 111.27** | 0.008** | 0.004** | 35.08** | 11.53** |
| Replication (WT) | 6 | 13.60 | 12.55 | 0.43 E<sup>-3</sup> | 0.33 E<sup>-3</sup> | 5.02 | 3.68 | 0.42 E<sup>-3</sup> | 0.26 E<sup>-3</sup> | 0.70 | 0.53 |
| Genotype (G) | 20 | 45.60** | 21.64** | 0.23** | 0.167** | 102.71** | 63.95** | 0.003** | 0.14 E<sup>-2</sup> | 2.25** | 0.93** |
| G × WT | 40 | 14.75** | 6.27** | 0.02** | 0.015** | 6.13** | 7.10** | 0.6 E<sup>-3**</sup> | 0.2 E<sup>-3**</sup> | 0.60** | 0.23** |
| GCA | 5 | 61.74** | 40.78** | 0.59** | 0.553** | 219.99** | 182.17** | 0.007** | 0.004** | 2.51** | 1.64** |
| SCA | 15 | 40.22** | 15.26** | 0.11** | 0.038** | 63.61** | 24.54** | 0.002** | 0.6 E<sup>-3**</sup> | 2.16** | 0.69** |
| GCA × WT | 10 | 35.18** | 19.27** | 0.02** | 0.022** | 8.65** | 13.54** | 0.001** | 0.6 E<sup>-3**</sup> | 1.13** | 0.69** |
| SCA × WT | 30 | 7.94** | 1.94** | 0.01** | 0.012** | 5.29** | 4.95** | 0.4 E<sup>-3**</sup> | 0.1 E<sup>-3**</sup> | 0.42** | 0.08* |
| Error | 120 | 1.12 | 1.10 | 0.54 E<sup>-3</sup> | 0.87 E<sup>-3</sup> | 1.98 | 2.09 | 3.87 E<sup>-5</sup> | 3.1 E<sup>-5</sup> | 0.05 | 0.05 |

\*\* and \* are significant at 1% and 5% levels of probability, respectively. Fruit yield (FY), essential oil content (EOC), essential oil yield (EOY), fatty oil content (FOC), fatty oil yield (FOY).

Analysis of variance for genetic effects revealed that both additive and non-additive gene actions are involved in the expression of traits in both F_1_ hybrids and F_2_ generations. Also, significant GCA × environment and SCA × environment interactions effect for all traits in both F_1_ and F_2_ generations (Table 3) reveal that general combing ability of parents and specific combining ability of hybrids were differently determined by additive and non-additive gene actions under different water treatments, respectively. Therefore, selection for parent with high GCA or hybrid with high SCA should be done according to the condition of target cultivating environment.

### Effect of water deficit stress on measured traits

Generally, results indicated that fruit yield, essential oil yield, fatty oil content and fatty oil yield were negatively affected by water deficit stress in both F_1_ hybrids and F_2_ generations in coriander. But essential oil content was significantly increased under water deficit stress. (Table 4).

**Table 4.** The mean of traits under different irrigation treatments in F_1_ and F_2_ generations of coriander.

| Water treatment | FY |  | EOC |  | FOC |  | EOY |  | FOY |  |
| --- | --- | --- | --- | --- | --- | --- | --- | --- | --- | --- |
|  | F<sub>1</sub> | F<sub>2</sub> | F<sub>1</sub> | F<sub>2</sub> | F<sub>1</sub> | F<sub>2</sub> | F<sub>1</sub> | F<sub>2</sub> | F<sub>1</sub> | F<sub>2</sub> |
| Well-watered | 9.19<sup>a</sup> | 6.74<sup>a</sup> | 0.351<sup>c</sup> | 0.337<sup>c</sup> | 20.59<sup>a</sup> | 18.35<sup>a</sup> | 0.035<sup>a</sup> | 0.023<sup>a</sup> | 1.88<sup>a</sup> | 1.22<sup>a</sup> |
| Moderate water Stressed | 4.51<sup>b</sup> | 3.94<sup>b</sup> | 0.530<sup>a</sup> | 0.446<sup>a</sup> | 18.60<sup>b</sup> | 17.76<sup>b</sup> | 0.029<sup>b</sup> | 0.021<sup>a</sup> | 0.87<sup>b</sup> | 0.73<sup>b</sup> |
| Severe water Stressed | 2.35<sup>c</sup> | 2.18<sup>c</sup> | 0.477<sup>b</sup> | 0.377<sup>b</sup> | 16.83<sup>c</sup> | 15.81<sup>c</sup> | 0.013<sup>c</sup> | 0.009<sup>b</sup> | 0.43<sup>c</sup> | 0.37<sup>c</sup> |

In each column the values with common letters do not differ significantly. Fruit yield (FY), essential oil content (EOC), essential oil yield (EOY), fatty oil content (FOC), fatty oil yield (FOY).

### Effect of water deficit stress on fruit yield

As shown in table 4, fruit yield was significantly affected by water treatments. The highest fruit yield obtained in well-watered condition while the minimum fruit yield obtained in severe water deficit stress in both F_1_ hybrids and F_2_ generations. A reduction in fruit yield of coriander under water deficit condition also reported by Nadjafi et al. (2009) and Khodadadi et al. (2016b). In other aromatic and medicinal crops, similar results observed by Zehtab-Salmasi et al. (2006) in dill (*Anethum graveolens* L.), Bannayan et al. (2008) in Plantago ovata and Nigella sativa, Laribi et al. (2009) in caraway (*Carum carvi* L.), Ekren et al. (2012) in purple basil (*Ocimum basilicum* L.) and Alinian and Razmjoo (2014) in cumin under drought stress condition. A fruit yield reduction under drought stress occurred through insufficient photosynthesis due to stomata closure and thereafter a reduction in CO_2_ uptake (Rebey et al., 2012), shortening flowering and fruit setting periods and preferential allocation of assimilates to the roots rather than the shoots (Alinian and Razmjoo, 2014).

### Effect of water deficit stress on essential oil content and essential oil yield

The largest value of essential oil content obtained in the moderate water deficit stress while the lowest essential oil content recorded in well-watered for both F_1_ hybrids and F_2_ generations. Results indicate that drought stress has a positive effect on the essential oil content in coriander. Increasing in the essential oil content by progress in drought stress has also been documented by Baher et al. (2002) in *Satureja hortensis* L, Yassen et al. (2003) in *Ocimum basilicum* L., Omidbaigi et al. (2003) in sweet basil, Dunford and Vazquez (2005) in Mexican oregano, Khalid (2006) in *Ocimum basilicum* L. and *Ocimum americanum* L., Petropoulos et al. (2008) in parsley, Bettaiebet al. (2009) in *Salvia officinalis* L., Ekren et al. (2012) in *Ocimum basilicum* L. and Alinian et al. (2014) in cumin.

Whereas, drought stress leads to decrease in essential oil yield in both F_1_ hybrids and F_2_ generations (Table 4). So that the highest value of essential oil yield obtained in the well-watered condition and the lowest essential oil yield observed in severe water deficit stress for both F_1_ hybrids and F_2_ generations (Table 4). Similar results were reported by Singh and Ramesh (2000), Zehtab-salmasi et al. (2001), Farahani et al. (2009) and Alinian and Razmjoo (2014). Essential oil yield depends on essential oil content and fruit yield. Because drought stress had a more reducing effect on fruit yield rather than an increasing effect on essential oil content, therefore, essential oil yield reduced under water deficit stress conditions (Farahani et al., 2009).

### Effect of water deficit stress on fatty oil content and yield

The largest fatty oil content and yield values obtained in well-watered and the least fatty oil content and fatty oil yield values were obtained in severe water deficit stress for both F_1_ hybrids and F_2_ populations. Similarly, Singh and Ramesh (2000) in rosemary, Zehtab-Salmasi et al. (2006) in dill (*Anethum graveolens* L.), Hamrouni et al. (2001) in safflower, Bettaieb et al. (2009) in *Salvia officinalis* L. and Bettaieb et al. (2011) in cumin (*Cuminum cyminum* L.) observed that the significant decreasing effect of water deficit stress on fatty oil content and fatty oil yield.

### Nature of gene action

A significant GCA and SCA variances for all traits in both F_1_ hybrids and F_2_ populations indicate that both additive and non-additive gene actions are contributed to determine these traits. Khodadadi et al. (2016b) reported that both non-additive and additive gene actions for the inheritance of different traits are important in coriander.

GCA/SCA ratio reflects the degree of trait which transmitted to the progeny. When the GCA/SCA ratio are closer to unit and zero show that additive and non-additive gene actions are mostly involved in inheritance of the trait, respectively. Consideration the GCA/SCA ratio, non-additive gene action was predominant for fruit yield, essential oil yield and fatty oil yield traits in F_1_ and F_2_ generations under well-watered condition (Table 5). The same gene action in F_1_ and F_2_ may be because of coupling phase linkage (Ramachandram and Goud, 1981). In advanced generations, when a coupling linkage present, additive genetic variance decrease and when the repulsion linkage present, additive genetic variance increase Robinson et al. (1960). Therefore, to improve fruit yield, essential oil yield and fatty oil yield traits under well-watered condition, selection should be delayed to the later generations of segregation. For fatty oil content, non-additive gene action nature was predominant in F_1_ hybrids, while in F_2_ generations the additive genetic effects were more important under well-watered condition (Table 5). The inconsistency in F_1_ and F_2_ results is due to the breakdown of dominance effects and gen linkages. Also, essential oil content was predominantly governed by additive gene action in both F_1_ hybrids and F_2_ generations. Presence of mostly additive gene action in F_2_ generation for fatty oil content and in both F_1_ and F_2_ generations for essential oil content suggests that selection programs can be effective in the F_2_ and later generations for improvement of fatty oil content and essential oil content traits under well-watered conditions.

**Table 5.** Analysis of variance for combining ability, variance components and GCA/SCA ratio.

| Water treatment | Estimate | FY |  | EOC |  | FOC |  | EOY |  | FOY |  |
| --- | --- | --- | --- | --- | --- | --- | --- | --- | --- | --- | --- |
|  |  | F<sub>1</sub> | F<sub>2</sub> | F<sub>1</sub> | F<sub>2</sub> | F<sub>1</sub> | F<sub>2</sub> | F<sub>1</sub> | F<sub>2</sub> | F<sub>1</sub> | F<sub>2</sub> |
| Well Watered | GCA | 31.82** | 13.86** | 0.131** | 0.128** | 59.34** | 30.62** | 0.002** | 0.001** | 16.25** | 8.30** |
|  | SCA | 21.19** | 4.85** | 0.018** | 0.014** | 28.44** | 6.88** | 0.001** | 0.26 E-3** | 26.53** | 6.08** |
|  | Error | 1.65 | 1.42 | 0.45 E-3 | 0.41 E-3 | 2.33 | 2.19 | 3.4 E-5 | 2.24 E-5 | 0.08 | 0.05 |
| | $\sigma_g^2$ | 2.21<sup>ns</sup> | 0.53* | 0.005** | 0.005** | 1.29<sup>ns</sup> | 0.99** | 4.5 E-5<sup>ns</sup> | 3.64 E-5** | 0.03<sup>ns</sup> | 0.004<sup>ns</sup> |
| | $\sigma_s^2$ | 18.21** | 1.83** | 0.006** | 0.004** | 8.70** | 1.56** | 0.4 E-3** | 7.96 E-5** | 0.64** | 0.08** |
|  | GCA/SCA | 0.12 | 0.37 | 0.62 | 0.68 | 0.23 | 0.56 | 0.18 | 0.48 | 0.09 | 0.10 |
| Moderate Water Stress | GCA | 65.85** | 48.31** | 0.323** | 0.307** | 101.93** | 119.15** | 0.006** | 0.003** | 2.873** | 2.147** |
|  | SCA | 16.30** | 8.64** | 0.074** | 0.041** | 23.03** | 16.00** | 0.001** | 5.3 E-4** | 0.791** | 0.448** |
|  | Error | 0.90 | 1.14 | 0.001 | 0.001 | 1.68 | 1.70 | 5.3 E-5 | 5.0 E-5 | 0.049 | 0.071 |
| | $\sigma_g^2$ | 2.06* | 1.65** | 0.010* | 0.011** | 3.29* | 4.30** | 1.8 E-4* | 1.2 E-4* | 0.006* | 0.003** |
| | $\sigma_s^2$ | 5.13** | 2.50** | 0.025** | 0.013** | 7.12** | 4.77** | 4.6 E-4** | 1.6 E-4** | 0.009** | 0.003** |
|  | GCA/SCA | 0.45 | 0.57 | 0.46 | 0.62 | 0.48 | 0.64 | 0.44 | 0.60 | 0.41 | 0.53 |
| Severe Water Stress | GCA | 13.62** | 11.30** | 0.177** | 0.161** | 76.03** | 59.48** | 6.4 E-4** | 3.9 E-4** | 0.68** | 0.48** |
|  | SCA | 4.75** | 3.58** | 0.044** | 0.008** | 22.73** | 11.56** | 2.3 E-4** | 8.4 E-5** | 0.20** | 0.12** |
|  | Error | 0.80 | 0.75 | 0.001 | 0.001 | 1.94 | 2.37 | 2.9 E-5 | 2.1 E-5 | 0.03 | 0.03 |
| | $\sigma_g^2$ | 0.37* | 0.32* | 0.006* | 0.006** | 2.22* | 2.00** | 1.7 E-5* | 1.3 E-5** | 0.02* | 0.02** |
| | $\sigma_s^2$ | 1.32** | 0.94** | 0.014** | 0.002** | 6.93** | 3.06** | 6.6 E-5** | 2.1 E-5** | 0.06** | 0.03** |
|  | GCA/SCA | 0.36 | 0.40 | 0.44 | 0.86 | 0.39 | 0.57 | 0.35 | 0.55 | 0.41 | 0.57 |

\*\* , \* and <sup>ns</sup> are significant at 1% and 5% level of probability and not significant, respectively. General combining ability (GCA), specific combining ability (SCA), fruit yield (FY), essential oil content (EOC), essential oil yield (EOY), fatty oil content (FOC), fatty oil yield (FOY).

In severe water deficit stress, results of GCA/SCA ratio for fruit yield showed that non-additive type of gene action was predominant in both F_1_ hybrids and F_2_ populations (Table 5). Therefore, to improve fruit yield under severe water deficit stress condition, selection should be delayed to the later generations of segregation to loss of non-additive gene actions. For fruit yield under moderate water deficit stress and essential oil content, fatty oil content, essential oil yield and fatty oil yield under both moderate and severe water deficit stress conditions, the non-additive gene action in F_1_ hybrids while an additive gene action in F_2_ generation were more important (Table 5). Therefore, breeding programs based on selection can be effective in the F_2_ and later generations for improvement of these traits under water deficit stress.

### Mean performance, heterosis and inbreeding depression Fruit yield

In well-watered condition, fruit yield varied from 2.40 (P_6_) to 9.71 g (P_2_) between the parents and ranged from 5.26 to 18.10 g (H_2_×_4_) between the F_1_ hybrids (Fig. 1A). Parental genotypes of the H_2_×_4_ had approximately half yield (6.80–9.71 g) as compared to their hybrid. In F_2_ generation, the fruit yield varied from 3.75 to 10.71 g between the hybrids (Fig. 1A). Similar to F_1_ generation, in F_2_ the highest fruit yield obtained by H_2_×_4_. Also, in F_1_ generations, almost all hybrids exhibited positive heterosis (7.82–115.40 %) in which P_4_ involved hybrids mostly showed high heterosis (+80.91 to +89.74 %). Inbreeding depression from F_1_ hybrids to F_2_ generations ranged from −7.94 % to −42.80 % for fruit yield (Fig. 1A).

**Fig. 1.**
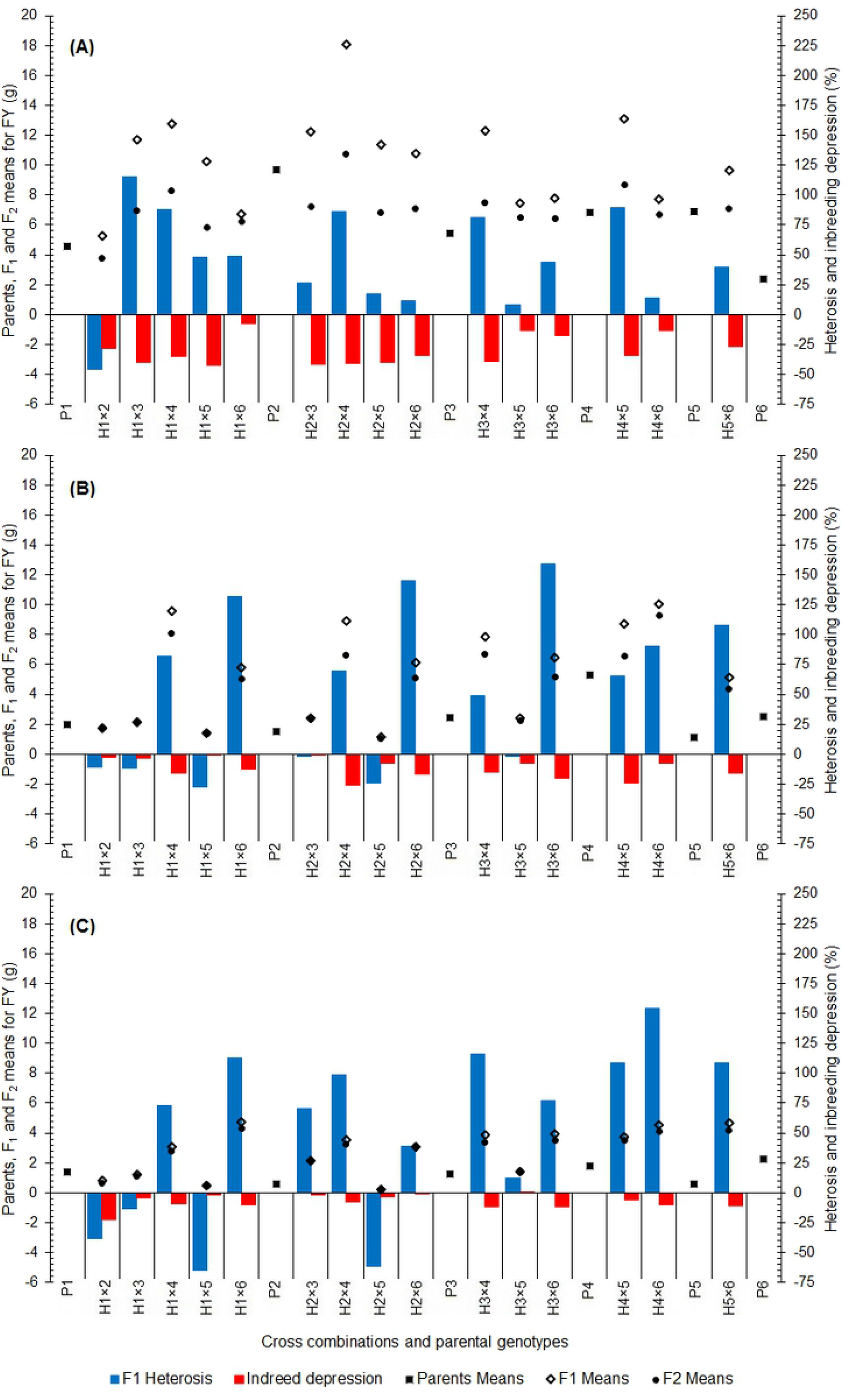
Mean, heterosis and inbreeding depression for fruit yield in F_1_ and F_2_ generations of coriander crosses. **A:** Well Watered, **B:** Moderate Water Stress, **C:** Severe Water Stress

In moderate water deficit stress condition, fruit yield varied from 1.14 (P_5_) to 5.27 g (P_4_) between the parents and ranged from 1.17 to 10.03 g between the F_1_ hybrids (Fig. 1B). A large fruit yield obtained in five F_1_ hybrids including H_4_×_6_ (10.03 g), H_1_×_4_ (9.58 g), H_2_×_4_ (8.93 g), H_4_×_5_ (8.71 g) and H_3_×_4_ (8.85 g). In F_2_ generation, fruit yield varied from 1.08 to 9.29 g (Fig. 1B). F_2_ generations relevant to the high yielding F_1_ hybrids also exhibited the highest fruit yield. When P_4_ and P_6_ contributed as one of the mating partners, the large heterosis vigor obtained (+107.40 % to +159.59 %). Inbreeding depression from F_1_ hybrids to F_2_ populations had larger range for fruit yield (−0.36 % to −26.05 %) in moderate water deficit stress than well-watered (Fig. 1B).

In severe water deficit stress, fruit yield varied from 0.58 (P_5_) to 2.24 g (P_6_) between parents and from 0.22 to 4.77 g between F_1_ hybrids (Fig. 1C). In F_2_ generation, fruit yield varied from 0.21 to 4.28 g (Fig. 1C) and a large fruit yield obtained from F_2_ populations derived from the P_4_ and P_6_ contributed hybrids. The heterosis values for fruit yield ranged between −64.68 and +154.54 % (Fig. 1C) and many of the hybrids exposed positive heterosis. Similar to moderate water deficit stress, inbreeding depression from F_1_ hybrids to F_2_ populations in severe water stress showed larger range (−0.59 to −22.66 %) than well-watered (Fig. 1C).

Higher heterosis and lower inbreeding depression in water deficit stressed conditions than those in well-watered condition reveal that the respective parents of hybrids probably were carriers of drought tolerance alleles could be homozygous recessive (Musembi et al., 2015). Therefore, their hybrids appeared superior in water deficit stressed conditions compared with the high yielding hybrids being superior in well water. In case of inbreeding depression from F_1_ hybrids to F_2_ generations, the heterozygote loci can maximally be 50 % breakdown. Therefore, an appearance of drought tolerance in F_2_ generations could yet be kept by heterozygote genes.

### Essential oil content

In well-watered treatment, the essential oil content ranged from 0.140 % (P_2_) to 0.550 % (P_4_) between the parents and from 0.250 to 0.563 % between the F_1_ hybrids (Fig. 2A). The highest essential oil content obtained in five hybrids of P_4_ (0.440–0.563 %), followed by H_1_×_3_ hybrid. In F_2_ generation, essential oil content ranged from 0.237 to 0.545% (Fig. 2A) and five of the F_2_ populations that a P_4_ was one of mating partner exposed the highest essential oil content (0.431– 0.545 %). In F_1_ generation (Fig. 2A) many of hybrids showed positive heterosis (+2.42 to +62.20 %). Also, all the F_2_ populations showed inbreeding depression (−2.07 to −9.06 %) (Fig. 2A).

**Fig. 2.**
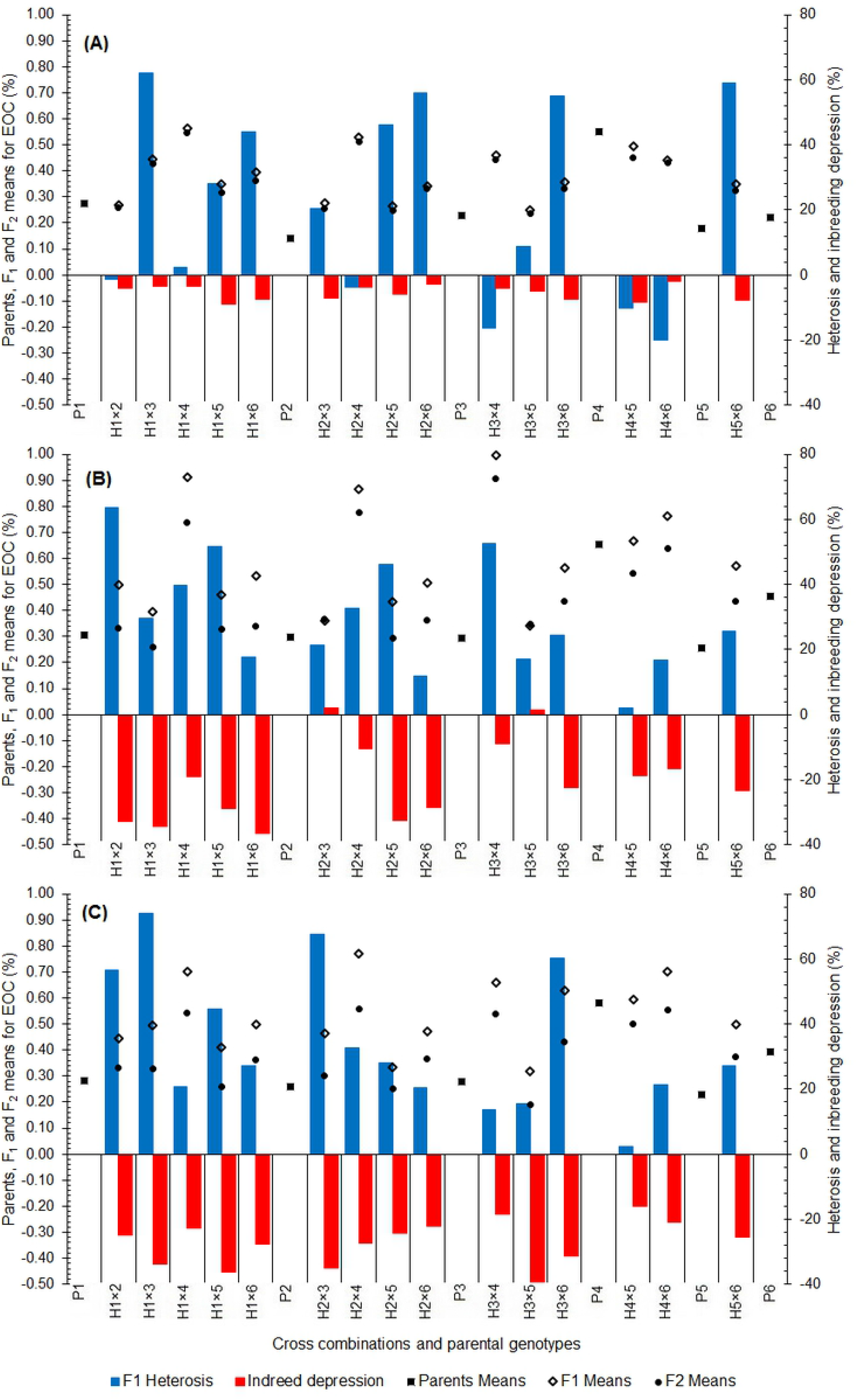
Mean, heterosis and inbreeding depression for essential oil content in F_1_ and F_2_ generations of coriander crosses. **A:** Well Watered, **B:** Moderate Water Stress, **C:** Severe Water Stress

In moderate water deficit stress, the essential oil content ranged from 0.257% (P_5_) to 0.653 % (P_4_) between the parents and from 0.343 to 0.997 % between the F_1_ hybrids (Fig. 2B). The highest essential oil content recorded in five hybrids relevant to P_4_ (0.667–0.997 %). In F_2_ generation, essential oil content ranged from 0.258 to 0.907 % between the populations (Fig. 2B) and similar to the F_1_ hybrids, five populations derived from P_4_ showed the highest essential oil content (0.542-0.907 %). In F_1_ generation all crosses exposed positive heterosis (+2.04 to +63.74 %) (Fig. 2B). Also, almost all the F_2_ populations showed inbreeding depression (−9.00 to −36.52 %) (Fig. 2B).

In severe water deficit stress, the essential oil content ranged from 0.227 % (P_5_) to 0.580 % (P_4_) between the parents and from 0.320 to 0.770 % between the F_1_ hybrids (Fig. 2C). The highest essential oil content obtained by five hybrids of P_4_ (0.593–0.770 %). In F_2_ generation, essential oil content was 0.191–0.560 % between the cross populations (Fig. 2C) and five derivatives of P_4_ showed high essential oil content (0.499–0.560 %). In F_1_ generation all hybrids showed positive heterosis (+2.30 to +74.12 %) and all of the F_2_ populations showed inbreeding depression (−15.89 to −40.38 %) (Fig. 2C).

The ranges of heterosis and inbreeding depression were higher in water deficit stressed conditions compare to the well water condition. Generally, high heterosis along with high inbreeding depression refers the presence of genes with non-additive action and high heterosis along with the least inbreeding depression indicates the presence of genes with additive action (Shukla and Gautam, 1990). Low inbreeding depression in well water condition suggests that increased vigor of F_1_s in such cases are expected to be mainly due to an accumulation of favorable additive action genes. Also, high inbreeding depression in water deficit stress condition indicates that non-additive action genes play major role in the inheritance of essential oil content. Our results are in accordance with previous researches on inbreeding depression under water deficit stressed conditions (Cheptou et al., 2000; Armbruster and Reed, 2005). In F_2_, even after inbreeding depression, some crosses exhibited good performance indicating the potential of these crosses to develop high essential oil content cultivars. The derivatives of the P_4_ parent displayed better mean performance as compared to their parents even after segregation and inbreeding depression. Therefore, P_4_ population could be used in the segregating generations to obtain genotypes with high essential oil content under different water treatments.

### Fatty oil content

In well-water, fatty oil content varied from 15.33 (P_4_) to 22 % (P_6_) between the parents and ranged from 16.33 to 26.67 % between the F_1_ hybrids (Fig. 3A). The highest fatty oil content recorded for hybrids of P_6_ (H_1_×_6_ (26.67 %), H_4_×_6_ (26.0 %), H_3_×_6_ (25.0 %) and H_2_×_6_ (23.0 %)) followed by H_1_×_4_ hybrid. Parental genotypes of these promising hybrids also had nearly high fatty oil content (18.33–22.0 %). In F_2_ generation, the fatty oil content varied from 14.94 to 22.54 % between the populations (Fig. 3A). The highest fatty oil content obtained in F_2_ generation by P_6_ hybrids and followed H_1_×_4_, H_2_×_5_, H_1_×_2_ hybrids. In F_1_ generation, heterosis ranged from +0.00 to +36.36 % for fatty oil content (Fig. 3A) and in F_2_ generation, inbreeding depression for fatty oil content observed from −8.32 to −25.75 % (Fig. 3A).

**Fig. 3.**
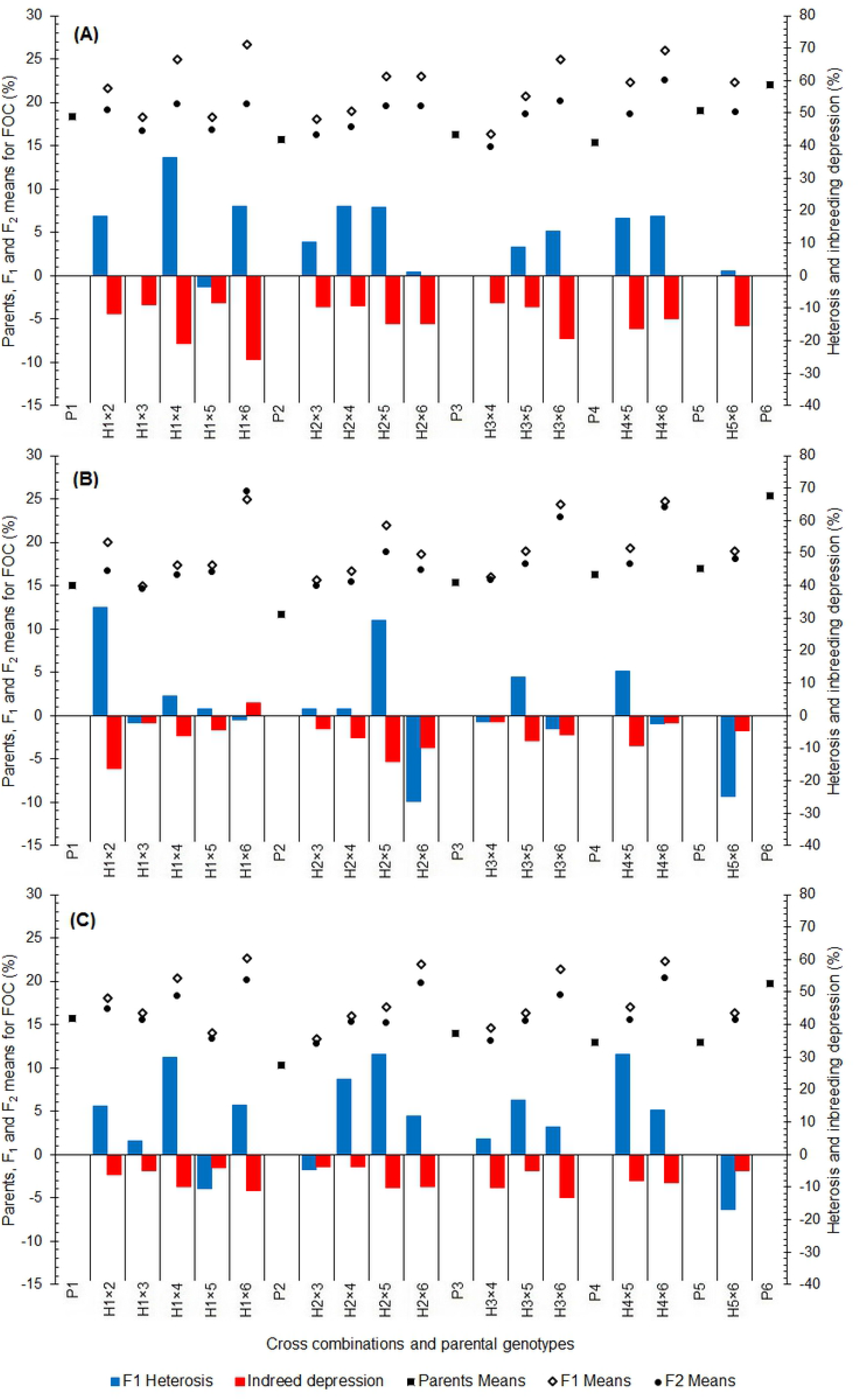
Mean, heterosis and inbreeding depression for fatty oil content in F_1_ and F_2_ generations of coriander crosses. **A:** Well Watered, **B:** Moderate Water Stress, **C:** Severe Water Stress

In moderate water deficit stress, the fatty oil content varied from 11.67 (P_2_) to 25.33 % (P_6_) and 15.00 to 25.0 % between parents and F_1_ hybrids, respectively (Fig. 3B). The highest fatty oil content observed in eight F_1_ hybrids that P_6_ involved in four crosses. In F_2_ generation, fatty oil content varied from 14.68 to 25.98 % between hybrids (Fig. 3B) and the highest fatty oil content (22.89–25.98 %) recorded for three hybrids of P_6_. The heterosis values for fatty oil content were +1.96 to +33.33 % (Fig. 3B) and almost all hybrids showed positive heterosis. F_2_ populations showed inbreeding depression for fatty oil content (−2.03 to −16.37 %) (Fig. 3B).

In severe water deficit stress, the fatty oil content varied from 10.33 (P_2_) to 19.67 % (P_6_) and 13.33 to 22.67 % between parents and F_1_ hybrids, respectively (Fig. 3C). The highest fatty oil content were recorded in F_1_ hybrids involving P_6_ and followed by H_1_×_4_ hybrid. In F_2_ generation, fatty oil content varied from 12.85 to 20.41 % between the hybrids (Fig. 3C) and the highest fatty oil content was obtained from hybrids of P_6_. The heterosis values for fatty oil content ranged from +4.26 to +30.77 % (Fig. 3C) and many of hybrids showed positive heterosis. The F_2_ generations displayed inbreeding depression (−3.64 to −13.30 %) for fatty oil content (Fig. 3C). Overall, it was revealed that P_6_ involved F_2_ populations could be utilize for developing cultivars with high fatty oil content under different water treatments.

The ranges of heterosis and inbreeding depression were higher in well-watered than water stressed conditions. High heterosis is well-known to be a result of the effects of non-additive genes (Shalaby, 2013; Solieman et al., 2013; Singh et al., 2014). Therefore, the higher heterosis and inbreeding depression in well water condition suggest that non-additive gene actions were more predominant in well water condition compare to the water deficit stressed conditions. F_2_ progenies derived from P_6_ contributed hybrids showed better mean performance even after inbreeding depression than their parents indicating the presence of transgressive segregation for fatty oil content under different water treatments.

### Essential oil yield and fatty oil yield

In well-watered treatment, the essential oil yield ranged from 0.005 (P_6_) to 0.037 g (P_4_) among the parents and from 0.014 to 0.096 g between the F_1_ hybrids (Fig. 4A). High essential oil yield was obtained for four P_4_ crosses (0.057–0.096 g). In F_2_ generation, essential oil yield ranged from 0.010–0.055 g between the cross generations (Fig. 4A) and four crosses of P_4_ showed a high essential oil yield (0.033–0.055 g). In F_1_ generation (Fig. 4A) almost all crosses indicated positive heterosis for essential oil yield (+7.48 to +213.91 %). Also, all of the F_2_ populations showed inbreeding depression (−15.06 to −47.80 %) (Fig. 4A).

**Fig. 4.**
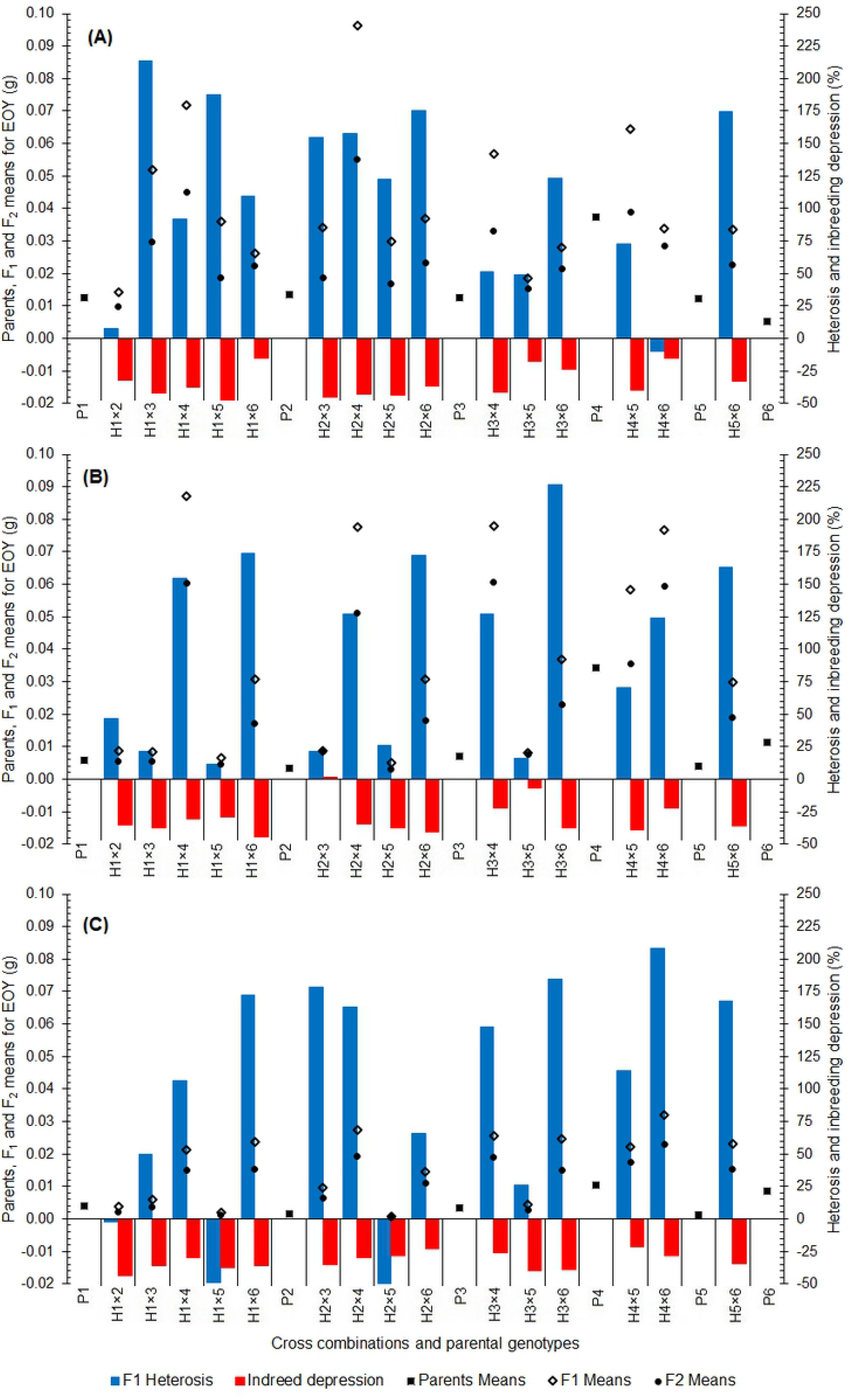
Mean, heterosis and inbreeding depression for essential oil yield in F_1_ and F_2_ generations of coriander crosses. **A:** Well Watered, **B:** Moderate Water Stress, **C:** Severe Water Stress

In moderate water stress, the essential oil yield ranged from 0.003 (P_2_) to 0.034 g (P_4_) between the parents and from 0.005 to 0.087 g between the F_1_ hybrids (Fig. 4B). Highest essential oil yield was recorded for five P_4_ crosses (0.058–0.087 g), followed by H_1_×_6_, H_3_×_6_, H_5_×_6_ hybrids. In F_2_ generation, essential oil yield ranged from 0.003–0.061 g between the cross population (Fig. 4B) and similar to the F_1_ generation, crosses of P_4_ showed highest essential oil yield (0.036–0.0.061 g). In F_1_ generation all crosses showed positive heterosis (+11.22 to +226.33 %) (Fig. 4B). Also, almost all of the F_2_ populations showed inbreeding depression for essential oil yield (−6.88 to −44.40 %) (Fig. 4B).

In severe water stress, the essential oil yield ranged from 0.002 (P_5_) to 0.010 g (P_4_) between the parents and from 0.001 to 0.032 g between the F_1_ hybrids (Fig. 4C). The highest essential oil yield was obtained in crosses of P_4_ (0.021–0.032 g), followed by H_1_×_6_, H_3_×_6_, H_5_×_6_ hybrids. In F_2_ generation, essential oil yield ranged from 0.001–0.023 g between the cross generations (Fig. 4C) and progenies of P_4_ and P_6_ showed the highest essential oil yield. In F_1_ generation, almost all crosses displayed positive heterosis (+26.01 to +208.31 %) (Fig. 4C). The F_2_ generation showed inbreeding depression (−21.96 to −40.85 %) (Fig. 4C). Overall, results indicated that P_4_ population could be used in the segregating generations to obtain genotypes with essential oil yield potential under different water treatments.

In well-water, the fatty oil yield varied from 1.12 to 3.41 g between parents and F_1_ hybrids (Fig. 5A). The highest fatty oil yield was obtained from H_2_×_4_, H_1_×_4_ hybrids. In F_2_ generation, fatty oil yield varied from 0.71 to 1.82 g between the generations (Fig. 5A) and highest fatty oil yield was noticed in generations derived from the hybrids of P_4_. The heterosis values for fatty oil yield were ranged from −26.95 to +204.96 % (Fig. 5A) and all hybrids showed positive heterosis. F_2_ populations displayed inbreeding depression for fatty oil yield (−21.88 to −49.31 %) (Fig. 5A).

**Fig. 5.**
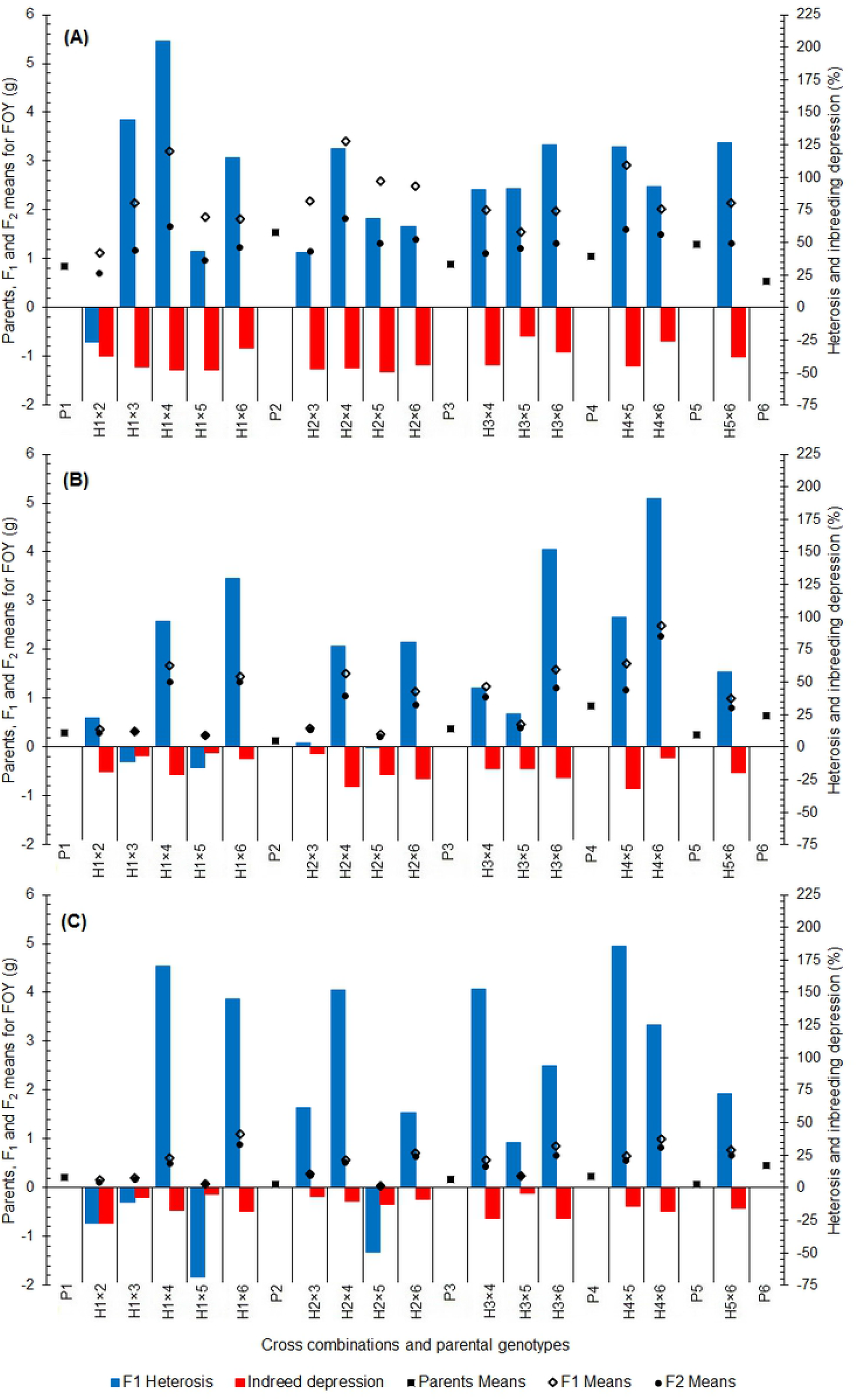
Mean, heterosis and inbreeding depression for fatty oil yield in F_1_ and F_2_ generations of coriander crosses. **A:** Well Watered, **B:** Moderate Water Stress, **C:** Severe Water Stress

In moderate water stress, the fatty oil yield ranged from 0.13 (P_2_) to 0.85 g (P_4_) between the parents and from 0.24 to 2.48 g between the F_1_ hybrids (Fig. 5B). High values of fatty oil yield were recorded in hybrids involving P_4_ and P_6_. In F_2_ generation, fatty oil yield ranged from 0.20– 0.2.27 g between the cross generations (Fig. 5B) and the crosses of P_4_ and P_6_ showed high fatty oil yield. In F_1_ generation (Fig. 5B) almost all of the hybrids showed positive heterosis (+3.42 to +191.18 %). Also, almost all of the F_2_ population showed inbreeding depression (−4.14 to −31.64 %) (Fig. 5B).

In severe water stress, the fatty oil yield varied from 0.06 (P_2_) to 0.45 g (P_6_) and 0.04 to 1.04 g between parents and F_1_ hybrids, respectively (Fig. 5C). High values of the fatty oil yield were recorded in F_1_ hybrids involving P_6_ and followed by hybrids of P_4_. In F_2_ generation, fatty oil yield varied from 0.03 to 0.89 g between the generations (Fig. 5C) and high values of the fatty oil yield was obtained from hybrids of P_6_. The heterosis values of fatty oil yield ranged from +35.04 to +185.27 % (Fig. 5C) and many of the hybrids showed positive heterosis. The F_2_ populations showed inbreeding depression (−4.53 to −27.02 %) (Fig. 5C). Overall, results indicated that P_6_ and P_4_ population could be used in the segregating generations to obtain genotypes with high fatty oil yield potential under different water treatments.

Inbreeding depression was higher in well water condition compare to water deficit stressed conditions for essential oil yield and fatty oil yield indicating that inbreeding depression was unstable across environments. Also, results revealed the higher heterosis values for essential oil yield and fatty oil yield than other traits indicating that non-additive genes were more responsible for the expression of these traits. These findings can be confirmed by the results of the GCA/SCA ratio in Table 5.

The utilization of hybrid vigor is one of the ways to improve yield in plant breeding. The existence of considerable degree of natural outcrossing had made these possible to use genetic diversity through production heterotic hybrids (Saxena *et al*., 1990). In coriander, heterosis cannot be exploited for higher production through commercial hybrids due to the nature of flower and poor seed recovery during hybridization. But estimation of heterosis for fruit yield, fatty oil and essential oils content will help in recognition crosses that can lead to isolate of advanced promising lines in segregating generation in coriander. Also, estimation of heterosis coupled with inbreeding depression shows that whether an amount of the vigor observed in segregating generations can be fixed in later generations by self-pollinating (Joseph and Santhoshkumar, 2000). The results showed that there was a positive heterosis for the traits examined in coriander which is an evidence for the existence of potential heterosis in Iranian coriander. In present study, the significant SCA effect indicates that there was non-additive gene effect, which could be the cause of the heterosis on the progenies observed and selection will not be effective in early generations. Hence, selection could be practiced in advance generations confirming to earlier reports.

The results showed that many of the F_2_ population exposed inbreeding depression and it was higher for fruit yield, essential oil yield and fatty oil yield. Inbreeding depression mostly was higher in hybrids with high performing than hybrids with low and moderate performing. Soomro and Kalhoro (2000), Khan et al. (2007) and Khan et al. (2009) reported that F_1_ hybrids with high performing were also correlated with higher inbreeding depression. Showing heterosis in F_1_ and inbreeding depression in F_2_ reveal the nature of gene action involved in the expression of the vigor in F_1_ and depression in F_2_. In F_2_ generation, the offspring’s of the parental genotypes P_4_ and P_6_ displayed better mean performance as compared to their parents and the selection in these crosses can provide transgressive gene recombinants for studied traits. P_4_ and P_6_ crosses are required to be subjected to the pedigree/progeny selection directly for reaching to the high potential cultivars. Also, P_4_ and P_6_ parents can be used as source of elite parents for synthetic cultivars (Khan et al., 2007; Khan et al., 2009) in coriander.

## Conclusion

Results indicated that water deficit stress negatively affected the fruit yield, essential oil yield, fatty oil content and fatty oil yield of coriander in both F_1_ and F_2_ generations. On the contrary, water deficit stress significantly increased the essential oil content of the coriander. Analysis of variance for genetic combining ability indicate that mean square due to GCA and SCA for all traits were highly significant in both F_1_ and F_2_ generations. Revealing the importance of additive and non-additive genetic nature in the expression of all traits in both F_1_ and F_2_ generations. Under water deficit stress conditions, non-additive gene action was predominant for studied traits in F_1_, while additive gene effects were more important in F_2_ generations except for fruit yield under severe water deficit stress. These results indicate that selection programs can be effective in the F_2_ and later generations (F_3_ or F_4_) for improvement of the studied traits under water deficit stress conditions. Also, for improvement of fruit yield under severe water deficit stress, selection should be delayed to later generations (F_3_ or F_4_) of segregation for dissipation of non-additive gene action. There was a positive heterosis in coriander for all traits. In F_2_, even after inbreeding depression, some promising generations displayed good performance and selection in such crosses can provide a better base for future. The progenies of the P_4_ and P_6_ parents displayed better mean performance as compared to their parents and the selection in these crosses provided transgressive gene recombinants for studied traits. It is also indicated that combined performance of F_1_ hybrids and F_2_ populations could be an appropriate criterion to recognizing the most promising populations to be used either as F_2_ hybrids or as a resource population for further selection in advanced generations.

## Notes

### Competing Interest Statement

The authors have declared no competing interest.

## References

Alinian, S., Razmjoo, J., 2014. Phenological, yield, essential oil yield and oil content of cumin accessions as affected by irrigation regimes. Ind. Crop. Prod. 54, 167–174.

Alinian, S., Razmjoo, J., Zeinali, H., 2016. Flavonoids, anthocynins, phenolics and essential oil produced in cumin (Cuminum cyminum L.) accessions under different irrigation regimes. Ind. Crop. Prod. 81, 49–55.

Armbruster, P., Reed, D., 2005. Inbreeding depression in benign and stressful environments. Heredity 95, 235–242.

Baher, Z.F., Mirza, M., Ghorbanli, M., Bagher Rezaii, M., 2002. The influence of water stress on plant height, herbal and essential oil yield and composition in Satureja hortensis L. Flavour Fragr. J. 17, 275–277.

Baker, R., 1978. Issues in diallel analysis. Crop Sci. 18, 533–536.

Bannayan, M., Nadjafi, F., Azizi, M., Tabrizi, L., Rastgoo, M., 2008. Yield and seed quality of Plantago ovata and Nigella sativa under different irrigation treatments. Ind. Crop. Prod. 27, 11–16.

Begnami, A., Duarte, M., Furletti, V., Rehder, V., 2010. Antimicrobial potential of Coriandrum sativum L. against different Candida species in vitro. Food Chem. 118, 74–77.

Bettaieb, I., Knioua, S., Hamrouni, I., Limam, F., Marzouk, B., 2011. Water-deficit impact on fatty acid and essential oil composition and antioxidant activities of cumin (Cuminum cyminum L.) aerial parts. J. Agr. Food Chem. 59, 328–334.

Bettaieb, I., Zakhama, N., Wannes, W.A., Kchouk, M., Marzouk, B., 2009. Water deficit effects on Salvia officinalis fatty acids and essential oils composition. Sci. Hortic. 120, 271–275.

Blank, A.F., Santa Rosa, Y.R., de Carvalho Filho, J.L.S., dos Santos, C.A., de Fátima Arrigoni-Blank, M., dos Santos Niculau, E., Alves, P.B., 2012. A diallel study of yield components and essential oil constituents in basil (Ocimum basilicum L.). Ind. Crop. Prod. 38, 93–98.

Burt, S., 2004. Essential oils: their antibacterial properties and potential applications in foods—a review. Int. J. Food Microbiol. 94, 223–253.

Charles, D.J., Joly, R.J., Simon, J.E., 1990. Effects of osmotic stress on the essential oil content and composition of peppermint. Phytochemistry 29, 2837–2840.

Cheptou, P.O., Berger, A., Blanchard, A., Collin, C., Escarre, J., 2000. The effect of drought stress on inbreeding depression in four populations of the Mediterranean outcrossing plant Crepis sancta (Asteraceae). Heredity 85, 294–302.

Chithra, V., Leelamma, S., 2000. Coriandrum sativum—effect on lipid metabolism in 1, 2-dimethyl hydrazine induced colon cancer. J. Ethnopharmacol. 71, 457–463.

Donega, M.A., Mello, S.C., Moraes, R.M., Cantrell, C.L., 2013. Nutrient uptake, biomass yield and quantitative analysis of aliphatic aldehydes in cilantro plants. Ind. Crop. Prod. 44, 127–131.

Dunford, N.T., Vazquez, R.S., 2005. Effect of water stress on plant growth and thymol and carvacrol concentrations in Mexican oregano grown under controlled conditions. J. Appl. Hortic. 7, 20–22.

Ekren, S., Sönmez, Ç., Özçakal, E., Kurttaş, Y.S.K., Bayram, E., Gürgülü, H., 2012. The effect of different irrigation water levels on yield and quality characteristics of purple basil (Ocimum basilicum L.). Agri. water manage. 109, 155–161.

El-Gabry, M., Solieman, T., Abido, A., 2014. Combining ability and heritability of some tomato (Solanum lycopersicum L.) cultivars. Sci. Hortic. 167, 153–157.

Farahani, H.A., Valadabadi, S.A., Daneshian, J., Shiranirad, A.H., Khalvati, M.A., 2009. Medicinal and aromatic plants farming under drought conditions. J. Hortic. Forest. 1, 086–092.

Farooq, M., Wahid, A., Kobayashi, N., Fujita, D., Basra, S., 2009. Plant drought stress: effects, mechanisms and management. Agron. Sustain. Dev. 29, 185–212.

Fonseca, S., Patterson, F.L., 1968. Hybrid vigor in a seven-parent diallel cross in common winter wheat (Triticum aestivum L.). Crop Sci. 8, 85–88.

Gallagher, A., Flatt, P., Duffy, G., Abdel-Wahab, Y., 2003. The effects of traditional antidiabetic plants on in vitro glucose diffusion. Nut. Res. 23, 413–424.

Griffing, B., 1956. A generalized treatment of the use of diallel crosses in quantitative inheritance. Heredity 10, 31–50.

Hamrouni, I., Salah, H.B., Marzouk, B., 2001. Effects of water-deficit on lipids of safflower aerial parts. Phytochemistry 58, 277–280.

Jaafar, H.Z., Ibrahim, M.H., Mohamad Fakri, N.F., 2012. Impact of soil field water capacity on secondary metabolites, phenylalanine ammonia-lyase (PAL), maliondialdehyde (MDA) and photosynthetic responses of Malaysian Kacip Fatimah (Labisia pumila Benth). Molecules 17, 7305–7322.

Joseph, J., Santhoshkumar, A., 2000. Heterosis and inbreeding depression in green gram (Vigna radiata L. Wilczek). Legume Res. 23, 118–121.

Kaushik, P., Plazas, M., Prohens, J., Vilanova, S. and Gramazio, P. (2018). Diallel genetic analysis for multiple traits in eggplant and assessment of genetic distances for predicting hybrids performance. Plos one, 13: e0199943.

Khalid, K.A., 2006. Influence of water stress on growth, essential oil, and chemical composition of herbs (Ocimum sp.). Int. Agrophys. 20, 289–296.

Khan, N.U., Hassan, G., Kumbhar, M.B., Kang, S., Khan, I., Parveen, A., Aiman, U., 2007. Heterosis and inbreeding depression and mean performance in segregating generations in upland cotton. Eur. J. Sci. Res. 17, 531–546.

Khan, N.U., Hassan, G., Kumbhar, M.B., Marwat, K.B., Khan, M.A., Parveen, A., Saeed, M., 2009. Combining ability analysis to identify suitable parents for heterosis in seed cotton yield, its components and lint % in upland cotton. Ind. Crop. Prod. 29, 108–115.

Khodadadi, M., Dehghani, H., Jalali-Javaran, M., 2017. Quantitative genetic analysis reveals potential to genetically improve fruit yield and drought resistance simultaneously in coriander. Front. Plant Sci. 8, 568.

Khodadadi, M., Dehghani, H., Jalali-Javaran, M., Rashidi-Monfared, S., Christopher, J.T., 2016a. Numerical and graphical assessment of relationships between traits of the Iranian Coriandrum sativum L. core collection by considering genotype × irrigation interaction. Sci. Hortic. 200, 73–82.

Khodadadi, M., Dehghani, H., Javaran, M.J., Christopher, J.T., 2016b. Fruit yield, fatty and essential oils content genetics in coriander. Ind. Crop. Prod. 94, 72–81.

Laribi, B., Bettaieb, I., Kouki, K., Sahli, A., Mougou, A., Marzouk, B., 2009. Water deficit effects on caraway (Carum carvi L.) growth, essential oil and fatty acid composition. Ind. Crop. Prod. 30, 372–379.

Lo Cantore, P., Iacobellis, N.S., De Marco, A., Capasso, F., Senatore, F., 2004. Antibacterial activity of Coriandrum sativum L. and Foeniculum vulgare Miller var. vulgare (Miller) essential oils. J. Agri. Food Chem. 52, 7862–7866.

Lubbe, A., Verpoorte, R., 2011. Cultivation of medicinal and aromatic plants for specialty industrial materials. Ind. Crop. Prod. 34, 785–801.

Matasyoh, J., Maiyo, Z., Ngure, R., Chepkorir, R., 2009. Chemical composition and antimicrobial activity of the essential oil of Coriandrum sativum. Food Chem. 113, 526–529.

Msaada, K., Hosni, K., Taarit, M.B., Chahed, T., Hammami, M., Marzouk, B., 2009a. Changes in fatty acid composition of coriander (Coriandrum sativum L.) fruit during maturation. Ind. Crop. Prod. 29, 269–274.

Msaada, K., Hosni, K., Taarit, M.B., Chahed, T., Kchouk, M.E., Marzouk, B., 2007. Changes on essential oil composition of coriander (Coriandrum sativum L.) fruits during three stages of maturity. Food Chem. 102, 1131–1134.

Msaada, K., Taarit, M.B., Hosni, K., Hammami, M., Marzouk, B., 2009b. Regional and maturational effects on essential oils yields and composition of coriander (Coriandrum sativum L.) fruits. Sci. Hortic. 122, 116–124.

Murphy, D.J., 1996. Engineering oil production in rapeseed and other oil crops. Trends. Biotechnol. 14, 206–213.

Murphy, D.J., Richards, D., Taylor, R., Capdevielle, J., Guillemot, J.C., Grison, R., Fairbairn, D., Bowra, S., 1994. Manipulation of seed oil content to produce industrial crops. Ind. Crops. Prod. 3, 17–27.

Musembi, K.B., Githiri, S.M., Yencho, G.C., Sibiya, J., 2015. Combining ability and heterosis for yield and drought tolerance traits under managed drought stress in sweet potato. Euphytica 201, 423–440.

Nadjafi, F., Damghani, A.M., Ebrahimi, S.N., 2009. Effect of irrigation regimes on yield, yield components, content and composition of the essential oil of four Iranian land races of coriander (Coriandrum sativum). J. Essent. Oil Bear. Pl. 12, 300–309.

Neffati, M., Marzouk, B., 2008. Changes in essential oil and fatty acid composition in coriander (Coriandrum sativum L.) leaves under saline conditions. Ind. Crop. Prod. 28, 137–142.

Neffati, M., Marzouk, B., 2010. Salinity impact on growth, essential oil content and composition of coriander (Coriandrum sativum L.) stems and leaves. J. Essent. Oil Bear. Pl. 22, 29–34.

Neffati, M., Sriti, J., Hamdaoui, G., Kchouk, M.E., Marzouk, B., 2011. Salinity impact on fruit yield, essential oil composition and antioxidant activities of Coriandrum sativum fruit extracts. Food Chem. 124, 221–225.

Omidbaigi, R., Hassani, A., Sefidkon, F., 2003. Essential oil content and composition of sweet basil (Ocimum basilicum) at different irrigation regimes. J. Essent. Oil Bear. Pl. 6, 104–108.

Petropoulos, S., Daferera, D., Polissiou, M., Passam, H., 2008. The effect of water deficit stress on the growth, yield and composition of essential oils of parsley. Sci. Hortic. 115, 393–397.

Ramachandram, M., Goud, J., 1981. Genetic analysis of seed yield, oil content and their components in safflower (Carthamus tinctorius L.). Theor. Appl. Genet. 60, 191–195.

Ramadan, M., Morsel, J.T., 2002. Oil composition of coriander (Coriandrum sativum L.) fruit-seeds. Eur. Food Res. Technol. 215, 204–209.

Ramadan, M.F., Morsel, J.T., 2006. Screening of the antiradical action of vegetable oils. J. Food Compos. Anal. 19, 838–842.

Rebey, I.B., Jabri-Karoui, I., Hamrouni-Sellami, I., Bourgou, S., Limam, F., Marzouk, B., 2012. Effect of drought on the biochemical composition and antioxidant activities of cumin (Cuminum cyminum L.) seeds. Ind. Crop. Prod. 36, 238–245.

Rieseberg, L.H., Archer, M.A., Wayne, R.K., 1999. Transgressive segregation, adaptation and speciation. Heredity 83, 363–372.

Robinson, H., Cockerham, C.C., Moll, R., 1960. Studies on estimation of dominance variance and effects of linkage bias. Biometrical genetics. Pergamon Press, New York, 171–177.

SAS Institute Inc., 1992. SAS Technical Report. SAS statistics Software: Changes and Enhancements. Release 6.07. SAS Institute Inc., Cary, North Carolina.

Saxena, K., Singh, L., Gupta, M., 1990. Variation for natural out-crossing in pigeonpea. Euphytica 46, 143–148.

Schegoscheski Gerhardt, I. F., Teixeira do Amaral Junior, A., Ferreira Pena, G., Moreira Guimarães, L. J., de Lima, V. J., Vivas, M., Araújo Diniz Santos, P. H., Alves Ferreira, F. R., Mendonça Freitas, M. S. and Kamphorst, S. H. (2019). Genetic effects on the efficiency and responsiveness to phosphorus use in popcorn as estimated by diallel analysis. Plos one, 14: e0216980.

Shalaby, T. A., 2013. Mode of gene action, heterosis and inbreeding depression for yield and its components in tomato (Solanum lycopersicum L.). Sci. Hortic. 164, 540–543.

Shukla, A., Gautam, N., 1990. Heterosis and inbreeding depression in okra (Abelmoschus esculentus L. Moench.). Indian J. Hortic. 47, 85–88.

Singh, M., Ramesh, S., 2000. Effect of irrigation and nitrogen on herbage, oil yield and wateruse efficiency in rosemary grown under semi-arid tropical conditions. J. Med. Aromatic Plant Sci. 22, 659–662.

Singh, P., Cheema, D., Dhaliwal, M., Garg, N., 2014. Heterosis and combining ability for earliness, plant growth, yield and fruit attributes in hot pepper (Capsicum annuum L.) involving genetic and cytoplasmic-genetic male sterile lines. Sci. Hortic. 168, 175–188.

Solieman, T., El-Gabry, M., Abido, A., 2013. Heterosis, potence ratio and correlation of some important characters in tomato (Solanum lycopersicum L.). Sci. Hortic. 150, 25–30.

Soomro, A., Kalhoro, A., 2000. Hybrid vigor (F_1_) and inbreeding depression (F_2_) for some economic traits in crosses between glandless and glanded cotton. Pak. J. Biol. Sci. 3, 2013–2015.

Sriti, J., Talou, T., Wannes, W.A., Cerny, M., Marzouk, B., 2009. Essential oil, fatty acid and sterol composition of Tunisian coriander fruit different parts. J. Sci. Food. Agr. 89, 1659–1664.

Teodoro, L. P. R., Bhering, L. L., Gomes, B. E. L., Campos, C. N. S., Baio, F. H. R., Gava, R., da Silva Júnior, C. A. and Teodoro, P. E. (2019). Understanding the combining ability for physiological traits in soybean. Plos one, 14: e0226523.

Townsend, T., Segura, V., Chigeza, G., Penfield, T., Rae, A., Harvey, D., Bowles, D. and Graham, I. A. (2013). The use of combining ability analysis to identify elite parents for Artemisia annua F1 hybrid production. Plos one, 8: e61989.

Turtola, S., Manninen, A.-M., Rikala, R., Kainulainen, P., 2003. Drought stress alters the concentration of wood terpenoids in Scots pine and Norway spruce seedlings. J. Chem. Ecol. 29, 1981–1995.

Wangensteen, H., Samuelsen, A.B., Malterud, K.E., 2004. Antioxidant activity in extracts from coriander. Food Chem. 88, 293–297.

Wong, P.Y., Kitts, D.D., 2006. Studies on the dual antioxidant and antibacterial properties of parsley (Petroselinum crispum) and cilantro (Coriandrum sativum) extracts. Food Chem. 97, 505–515.

Yassen, M., Ram, P., Yadav, A., Singh, K., 2003. Response of Indian basil (Ocimum basilicum) to irrigation and nitrogen schedule in Central Uttar Pradesh. Ann. Plant Physiol. 17, 177–181.

Zehtab-Salmasi, S., Ghassemi-Golez, K., Moghbeli, S., 2006. Effect of sowing date and limited irrigation on the seed yield and quality of dill (Anethum graveolens L.). Turk. J. Agric. For. 30, 281–286.

Zhang, Y., Kang, M.S., Lamkey, K.R., 2005. DIALLEL-SAS05: a comprehensive program for Griffing’s and Gardner-Eberhart analyses. Agron. J. 97, 1097–1106.

